# Genome-wide study of the effect of blood collection tubes on the cell-free DNA methylome

**DOI:** 10.1101/2020.04.23.055293

**Authors:** Ruben Van Paemel, Andries De Koker, Christa Caggiano, Annelien Morlion, Pieter Mestdagh, Bram De Wilde, Jo Vandesompele, Katleen De Preter

## Abstract

**Background:** The methylation pattern of cfDNA, isolated from liquid biopsies, is gaining substantial interest for diagnosis and monitoring of diseases. We have evaluated the impact of type of blood collection tube and time delay between blood draw and plasma preparation on bisulfite-based cfDNA methylation profiling.

**Methods:** 15 tubes of blood were drawn from three healthy volunteer subjects (BD Vacutainer K2E EDTA spray tubes, Streck Cell-Free DNA BCT tubes, PAXgene Blood ccfDNA tubes, Roche Cell-Free DNA Collection tubes and Biomatrica LBgard blood tubes in triplicate). Samples were either immediately processed or stored at room temperature for 24 or 72 hours before plasma preparation. DNA fragment size was evaluated by capillary electrophoresis. Reduced representation bisulfite sequencing was performed on the cell-free DNA isolated from these plasma samples. We evaluated the impact of blood tube and time delay on several quality control metrics.

**Results:** All preservation tubes performed similar on the quality metrics that were evaluated. Furthermore, a considerable increase in cfDNA concentration and the fraction of it derived from NK cells was observed after a 72-hour time delay in EDTA tubes.

**Conclusion:** The methylation pattern of cfDNA is robust and reproducible in between the different preservation tubes. EDTA tubes processed as soon as possible, preferably within 24 hours, are the most cost effective. If immediate processing is not possible, preservation tubes are valid alternatives.

## Introduction

The analysis of cell-free tumor-derived DNA (ctDNA) isolated from liquid biopsies, including blood plasma, has emerged as a complementary assay for tumor tissue genomic profiling^1,2^. The methylation pattern of cell-free DNA is currently under investigation as a biomarker for the non-invasive diagnosis of both benign^3^ and malignant^4^ conditions. Several algorithms are developed to deconvolute the epigenetic signal estimating the proportion of cell types contributing to cell-free DNA makeup^3,5,6^

The current lack of standardization of the pre-analytical phase is hampering routine clinical implementation. Pre-analytical factors like storage conditions and processing time impact cfDNA concentrations^7,8^. Blood tubes containing preservation agents aimed at stabilization of white blood cells inhibiting them from releasing DNA into the plasma have been developed^9^ to increase the allowed processing delay.

It is unknown to what extent the preservation agents have an impact on the methylation pattern of cell-free DNA. For example, formaldehyde results in deamination of cytosines, thereby resulting in C to T transitions^10^. In addition, crosslinking reagents can also induce methyl bridges with cytosine and guanine through a covalent bond^11^. These bonds can interfere with bisulfite conversion efficiency required for methylation analysis. As the makeup of preservation agents are usually proprietary information, the expected theoretical impact on cfDNA is generally unknown.

Although several manufacturers claim that their tube does not impact cfDNA methylation status^12,13^, studies are conflicting. One study suggests that PAXgene Blood DNA tubes and Streck Cell-Free DNA BCT tubes are unsuitable for the analysis of methylated cfDNA, as assessed with methylation-specific PCR (MSP) for SEPT9^14^. Schröder and Steimer reported a significantly higher methylation percentage in whole blood after ten months of storage^15^. In contrast, Zavridou and colleagues^16^ concluded that plasma samples from preservation tubes remain suitable for cfDNA methylation analysis with methylation specific PCR (MSP) when stored at -80 °C.

The major drawback of using methylation-specific PCR to evaluate pre-analytical variables is that only a limited number of loci are analyzed. To our knowledge, no study has systematically evaluated the effect of various preservation agents and the timing of plasma preparation after blood draw on the genome-wide methylation pattern. To this end, we have collected 45 blood samples in 5 different blood collection tubes (4 preservation tubes and 1 EDTA tube) from 3 healthy volunteers and performed plasma preparation at 3 different time points after blood draw followed by reduced representation bisulfite sequencing on the resulting cell-free DNA (cf-RRBS)^17^.

## Methods

### Healthy volunteers and samples

Blood was drawn from three healthy individuals (males and females between 24 and 45 years old). 15 tubes were drawn per healthy volunteer: 3x BD Vacutainer K2E EDTA spray tubes (REF 367525), 3x Streck Cell-Free DNA BCT tubes (catalogue number 218962), 3x QIAGEN PAXgene Blood ccfDNA tubes (catalogue number 768115), 3x Roche Cell-Free DNA Collection tubes (catalogue number 07785674001) and 3x Biomatrica LBgard blood tubes (SKU 68021-001), resulting in 45 samples in total.

The order of blood draw and collection in different tubes was randomized per donor. Seven tubes were collected from one antecubital vein and 8 tubes were collected from the contralateral antecubital vein with the BD Vacutainer push button 21G, 7” tube with pre-attached holder tube butterfly needle system (REF 367355). The ethical committee of Ghent University Hospital approved the study (identifier 2017/1207) and written consent was obtained from all volunteers enrolled in this study. The tubes are abbreviated as “EDTA” (BD Vacutainer K2E EDTA spray tubes), “Streck” (Streck Cell-Free DNA BCT tubes), “Roche” (Roche Cell-Free DNA collection tubes), “PAXgene” (QIAGEN PAXgene Blood ccfDNA tubes) or “Biomatrica” (Biomatrica LBgard blood tubes). Descriptive statistics were performed with R v3.5.1 and are reported as median and 25^th^ (Q25) and 75^th^ percentile (Q75) values [interquartile range].

### Sample collection and processing

After blood draw, samples were either immediately processed to plasma (1 per tube type, per donor; 15 tubes in total), or incubated at room temperature for 24 hours (n = 15) or 72 hours (n = 15). Tubes that were incubated were shielded from direct sunlight on a lab bench in a climate-controlled environment. Plasma was prepared according to a previously published protocol developed at the Ghent University Hospital^18^ (EDTA and Streck, 10 min at 1600 rcf followed by 10 minutes at 16000 rcf or PAXgene, 15 min at 1900 rcf) or according to the manufacturers’ guidelines (Roche, 10 min at 1600 rcf and Biomatrica, 15 min at 1900 rcf followed by 15 min at 4500 rcf). Acceleration and deceleration were set to 2 across all centrifugations (Eppendorf centrifuge 5804, Eppendorf). Plasma was aliquoted per 1500 µL and stored at -80°C until further processing.

### Cell-free DNA extraction

cfDNA was extracted using the Maxwell RSC LV ccfDNA kit (Promega). Isolation of cfDNA was done starting from 1500 µL of plasma. DNA extraction was performed according to the manufacturer’s instructions. DNA was eluted in 75 μL of elution buffer (Promega).

### Cell-free DNA quality control

DNA concentration was measured using the Qubit high-sensitivity kit (Thermo Fisher Scientific). DNA concentration and size distribution was measured using the FEMTO Pulse Automated Pulsed-Field CE Instrument (Agilent) according to the manufacturer’s instructions (NGS Kit, FP-1101-0275).

### RRBS library construction

Library construction was performed according to the methods described by De Koker et al.^17^ with the following 3 modifications: [1] The complete eluate that was remaining after quality control (2 µL Qubit and 2 µL Femto PULSE) was used for library construction (= 71 µL). For this, samples were concentrated with a vacuum centrifuge (SpeedVac, Thermo Fischer Scientific, V-AQ program) at 35 °C and nuclease-free water was added to a volume of 11.1 μL. Unmethylated lambda phage DNA (0.005 ng, or 0.5 µL of a 0.01 ng/µL solution) was added to the eluate after the SpeedVac step. [2] Libraries prepared using the cf-RRBS protocol were cleaned by magnetic bead selection (AMPure XT beads – NEB) and eluted in 0.1X TE buffer. The libraries were visualized with the Fragment Analyzer (Agilent) and quantified using the Kapa library quantification kit for Illumina platforms (Kapa Biosystems). [3] Based on the concentration, the libraries were equimolarly pooled and were sequenced on a NovaSeq 6000 instrument with a NovaSeq SP kit (paired-end, 2×50 cycles), using 3% phiX and a loading concentration between 1.8 and 2.5 nM. A maximum of 15 samples were pooled in one sequencing run, and samples from different donors and tubes were mixed to avoid sequencing batch effects.

### Sequencing quality control and mapping

After sequencing, bcl files were demultiplexed using bcl2fastq v2.19.1.403. The raw fastq files were first quality checked with FastQC v0.11.5^19^. Adaptors were removed with Trim Galore v0.6.0 (with flags --three_prime_clip_R1 1 --three_prime_clip_R2 1 --clip_R1 3 -- clip_R2 3 to remove methylation bias at the three or five prime end due to the MspI restriction digest) and CutAdapt v1.16^20^. Processed fastq files were again quality checked with FastQC. Mapping to GRCh37 was done with Bismark v0.20.1^21^ with default parameters. Optical duplicates were removed with Picard v2.18.27 (http://broadinstitute.github.io/picard/), with parameters OPTICAL_DUPLICATE_PIXEL_DISTANCE=12000, REMOVE_SEQUENCING_DUPLICATES=true, TAGGING_POLICY=OpticalOnly,READ_NAME_REGEX=’[a-zA-Z0-9]+:[0-9]+:[a-zA-Z0-9]+:[0-9]+:([0-9]+):([0-9]+):([0-9]+)_[0-9]+:[a-zA-Z0-9]+:[0-9]+:[a-zA-Z0-9]+’). The bait and target regions were defined as the MspI regions between 20-200 bp in GRCh37 (see ‘NNLS deconvolution) and the number of reads in these regions were counted with Picard (hsmetrics module. Per-sample information is available in supplementary table 1. Figures were made with the R programming language (see “code availability”)

### Quality control metric development

In order to evaluate tube stability across time points, we determined 6 metrics per blood collection tube: [1] percentage of reads mapping within MspI regions, [2] evolution of relative DNA concentration (as assessed by the ratio of reads mapping to the human genome versus reads mapping to the lambda genome), [3] genome-wide CpG methylation percentage, [4] Total number of CpGs covered with at least 15 reads, [5] reproducibility of individual cytosine in CpG context calls by calculating the area-left-of-the-curve^22^ (ALC) and [6] the evolution of the immune cell content.

In order to not selectively favor samples that have been sequenced deeper compared to others, we down-sampled all libraries to 18 million read pairs with seqtk v1.3 (random seed = 18).

For metrics that are described in percentages (e.g. reads mapping within MspI regions, genome wide methylated CpG, reproducibility with ALC, and immune cell contribution) the absolute change (e.g. genome-wide CpG % at T24 minus genome-wide CpG % at T0) was calculated across all timepoints per donor (excluding the T24-T72 comparison), resulting in 6 absolute change values. For all other metrics (cell-free DNA concentration and number of CpG reaching 15 reads), we calculated the fold-change (e.g. across all timepoints per donor, resulting in 6 fold-change values (e.g. for donor 1: T0 vs T24, T0 vs T72, times three donors, illustrated in figure 1).

**Figure 1:**
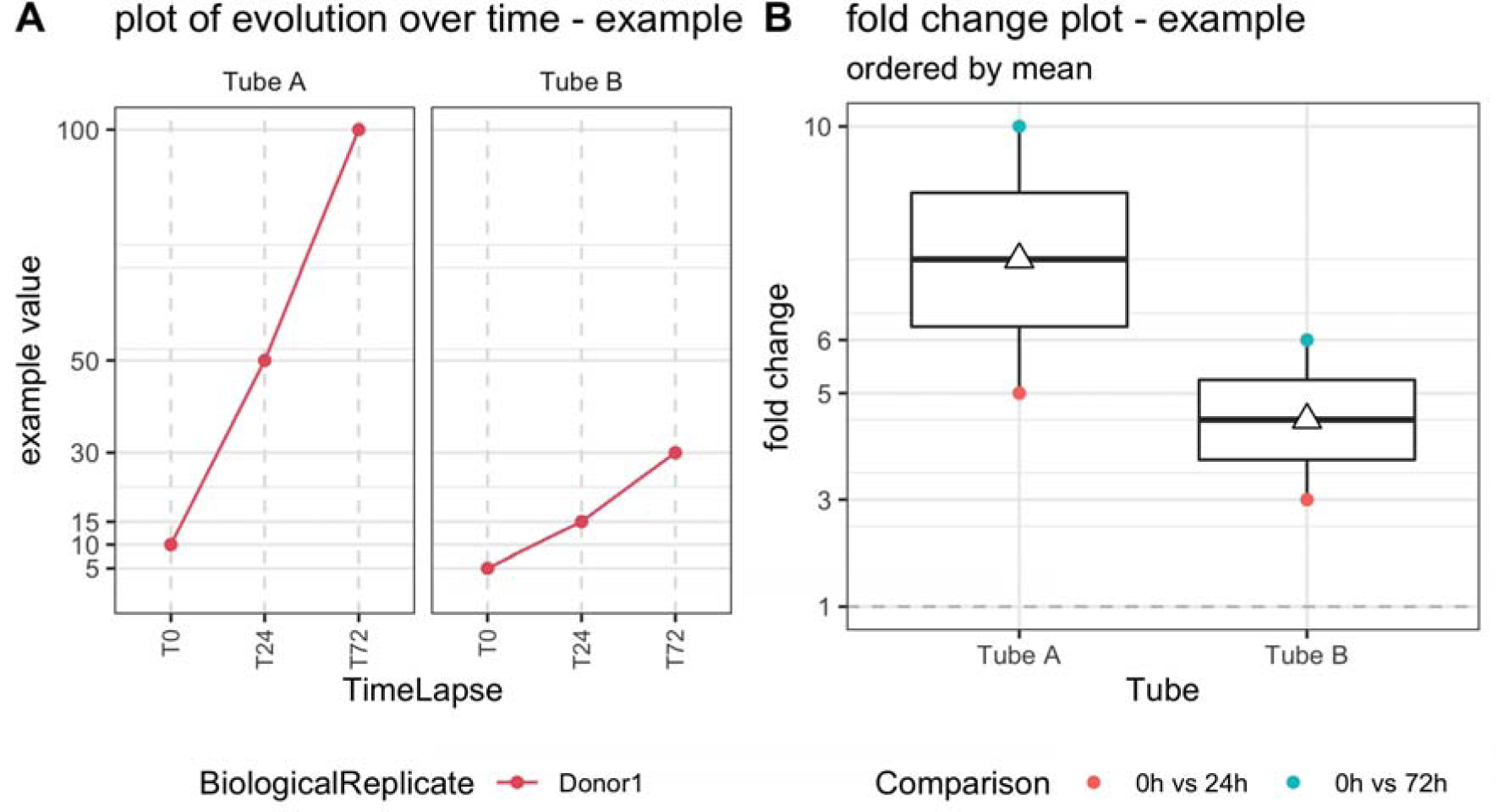
Illustrative example of quality control metric evolution over time for 1 donor, 2 tubes and 3 time points (left) and corresponding boxplot of the fold changes per tube (right). T0 = plasma prepared immediately after blood draw, T24 = plasma prepared 24 hours after blood draw, T72 = plasma prepared 72 hours after blood draw. The white triangle on the boxplot corresponds to the mean.

### Differential methylation of promotor regions

In order to assess differentially methylated promotor regions (DMRs), we used edgeR^23^ 3.28.1. Promotor regions were defined as 1000 bp upstream to 2000 bp downstream of transcription start sites (TSS). Uninformative regions (either completely methylated or completely unmethylated across all samples) and regions with fewer than 15 reads were removed from the dataset (n = 3983 remaining regions). Two types of comparisons were performed: (1) DMRs were determined by comparing all tubes with one another in a pairwise manner, exclusively at T0 and (2) DMRs were determined within one tube type by comparison T0 with T72. The false discovery rate (FDR) cut-off was specified at 0.05.

### NNLS deconvolution to determine cell type contribution

Deconvolution of plasma cfDNA samples was done with non-negative least squares (NNLS) matrix decomposition as described by Moss and colleagues3 (https://github.com/nloyfer/meth_atlas). CpGs were grouped in order to use more mappable reads. Input data (cf-RRBS data) was prepared similar to Kang et al.5 However, we adjusted these regions to make them more suited for cf-RRBS data. First, we used mkrrgenome^24^ to extract all MspI regions between 20-200 bp from GRCh37. Then, we merged all remaining regions within 1 bp from each other with BEDtools v2.27.1. Finally, clusters were retained if they contain at least 3 CpGs covered on the Illumina HM450K array, resulting in 14,103 clusters covering 61,750 probes on the HM450K array. We used the Illumina HM450K reference matrix as published by Moss and colleagues to perform the deconvolution. Similarly, we selected the top 100 most specific hypermethylated and hypomethylated regions per cell type as described by them.

### CelFiE deconvolution to determine cell type contribution

Deconvolution of plasma cfDNA samples was done with CelFiE (commit d3f975f) and its accompanying reference dataset^25^. In order to adapt the WGBS dataset (sourced from ENCODE^26^ and BLUEPRINT^27^) for cf-RRBS data, we removed all CpGs not covered in cf-RRBS from the WGBS dataset (based on the MspI regions between 20-200 bp generated with mkrrgenome). Afterwards, tissue informative markers were generated as described by Caggiano and colleagues^25^ and deconvolution was performed with CelFiE (https://github.com/christacaggiano/celfie), directly from coverage files obtained from Bismark.

## Results

### cfDNA quality control

Using the Qubit High Sensitivity kit, we were unable to determine the cfDNA concentration in each sample (n = 20 samples with a concentration below the Qubit quantitative range). Using the Femto PULSE, cfDNA concentrations could be assessed in all samples (3.6 ng/mL plasma [1.75-4.88]) (see next paragraph). At later timepoints, the second and third nucleosome peaks are less apparent on the capillary electropherograms in the Roche, Biomatrica and DNA Streck tubes (figure 2). cf-RRBS library construction was successful in all samples. After sequencing, all libraries were down-sampled to 18M read pairs. 95.0% [92.0 – 96.0] of the total reads were remaining after adaptor removal. One healthy volunteer showed fewer reads remaining after adaptor removal in all 15 samples (supplementary figure S1). Overall, 63.66% [62.92-64.67] of all reads mapped uniquely to the human reference genome (GRCh37, supplementary figure S2). Furthermore, we noticed a monotonous increase in mapping efficiency in EDTA tubes over time. We assessed bisulfite conversion efficiency by calculating the genome wide CHH methylation percentage. A conversion efficiency of 99.3% [99.2-99.4] was observed and was similar in all tubes and across all three time points (supplementary figure S4). We assessed MspI efficiency by evaluating the percentage of reads mapping within all theoretical MspI fragments between 20 and 200 bp. Overall, 89.71% [88.28-91.01] of reads mapped within these regions, with no tube showing a substantial and consistent change between time points (supplementary figure S5).

**Figure 2:**
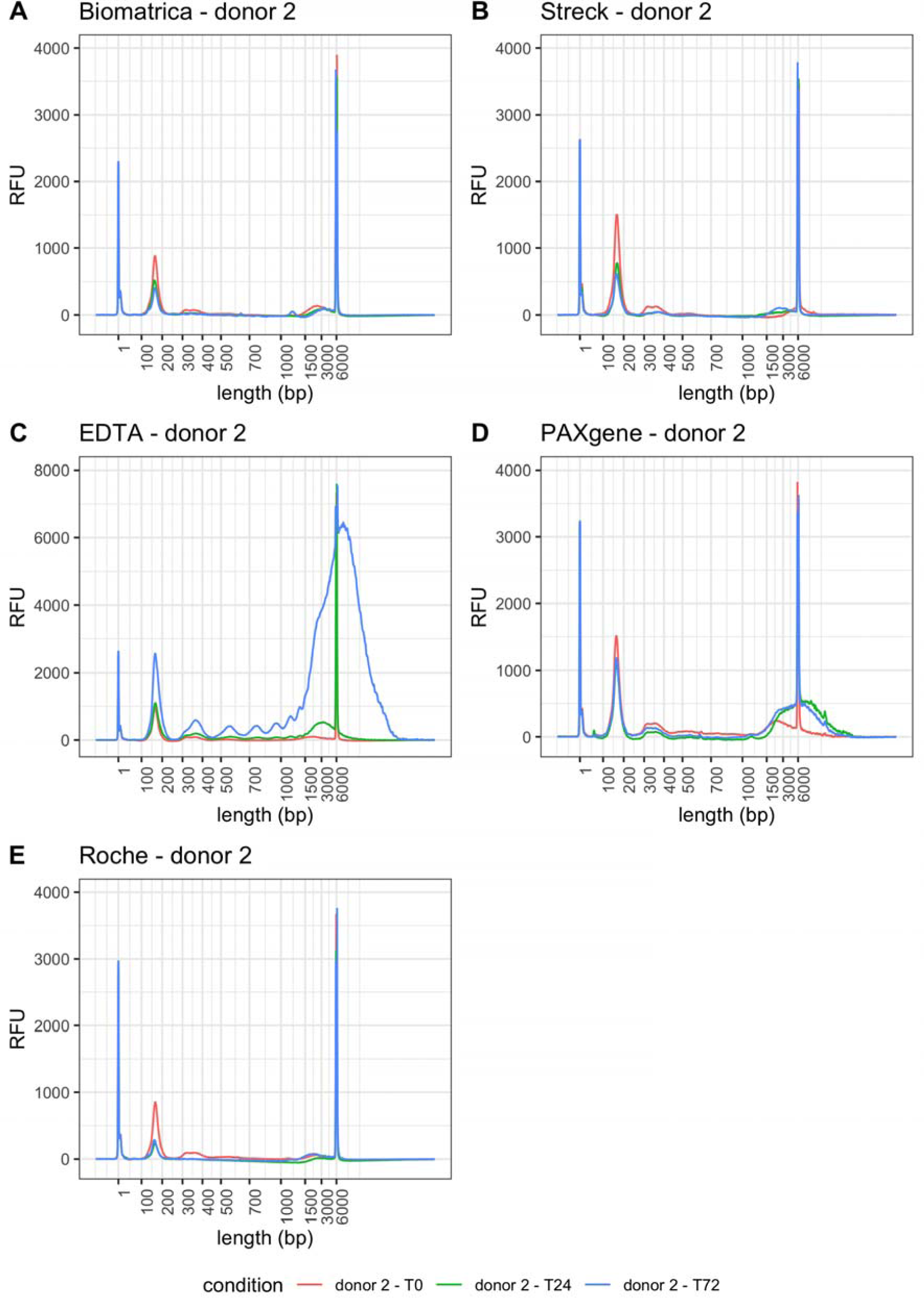
Capillary electropherograms (Femto PULSE) of all 15 cfDNA samples obtained from donor 2. RFU = relative fluorescence units. T0 = plasma prepared immediately after blood draw, T24 = plasma prepared 24 hours after blood draw, T72 = plasma prepared 72 hours after blood draw. Peaks at 1 and 6000 bp correspond to the lower marker (1 bp) and upper marker (6000 bp).

### cfDNA content

After DNA isolation and before library construction, 50 pg of lambda DNA was added to all samples. By assessing the ratio of the reads mapping to the human genome versus the reads mapping to the lambda genome, we calculated the (lambda equivalent) cfDNA concentration in ng/mL plasma (figure 3A & 3B). The EDTA tubes revealed a substantial increase in DNA concentration after 72 hours with an overall mean fold change of 4.06 (median concentration at T0 = 2.98 ng/mL plasma, median concentration at T72 = 12.65 ng/mL plasma). In contrast, the Roche tubes revealed a considerable decrease in DNA concentration after 72 hours with an overall mean fold change of 4.61 (median concentration at T0 = 2.45 ng/mL plasma, median concentration at T72 = 0.46 ng/mL plasma). We confirmed these findings by Femto PULSE quantification (figure 2, supplementary figures S13, S14). The cfDNA concentration measured with Femto PULSE correlated well with the sequencing based cfDNA (lambda equivalent) concentration determination (Pearson’s r = 0.73, supplementary figure S6).

**Figure 3:**
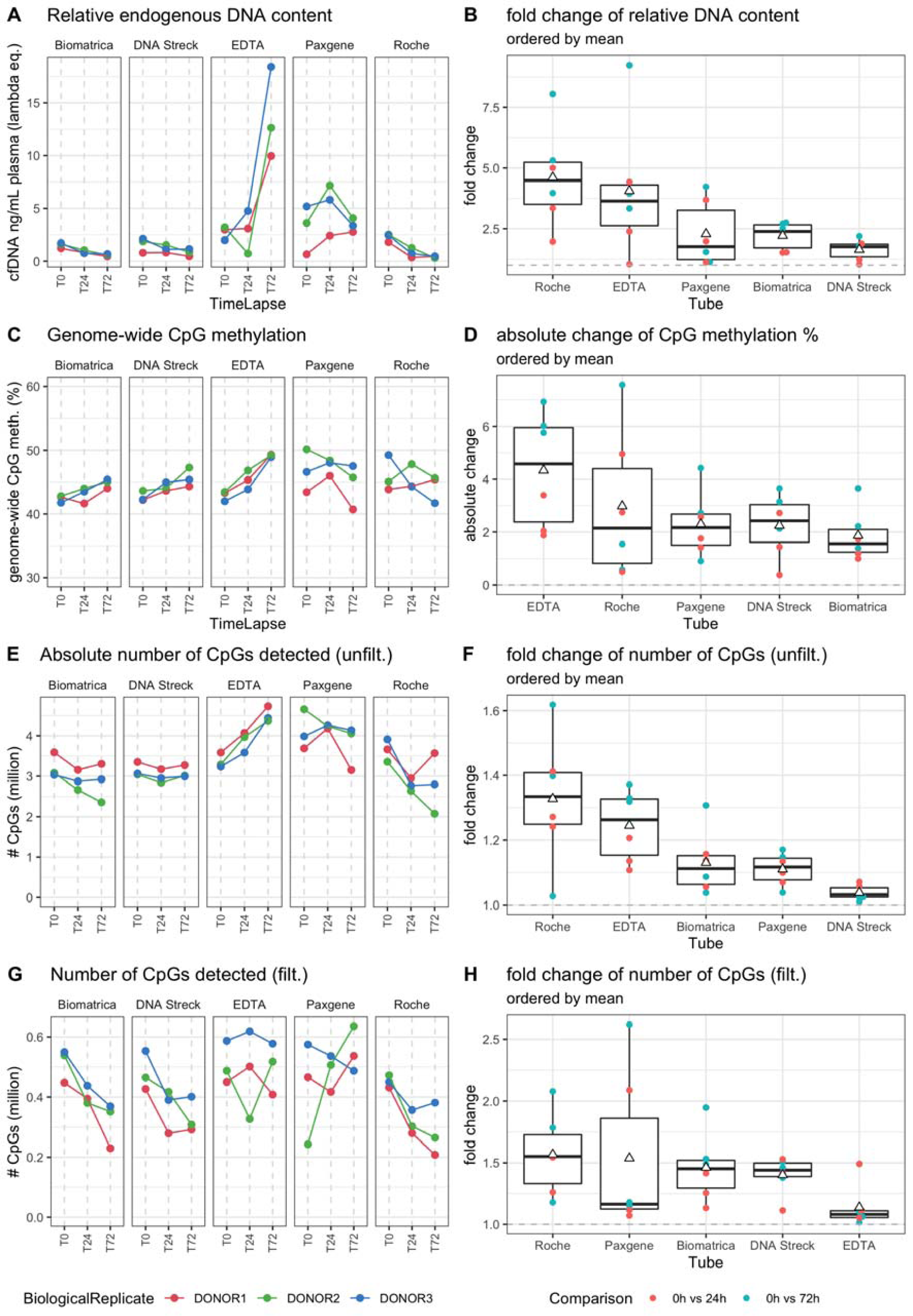
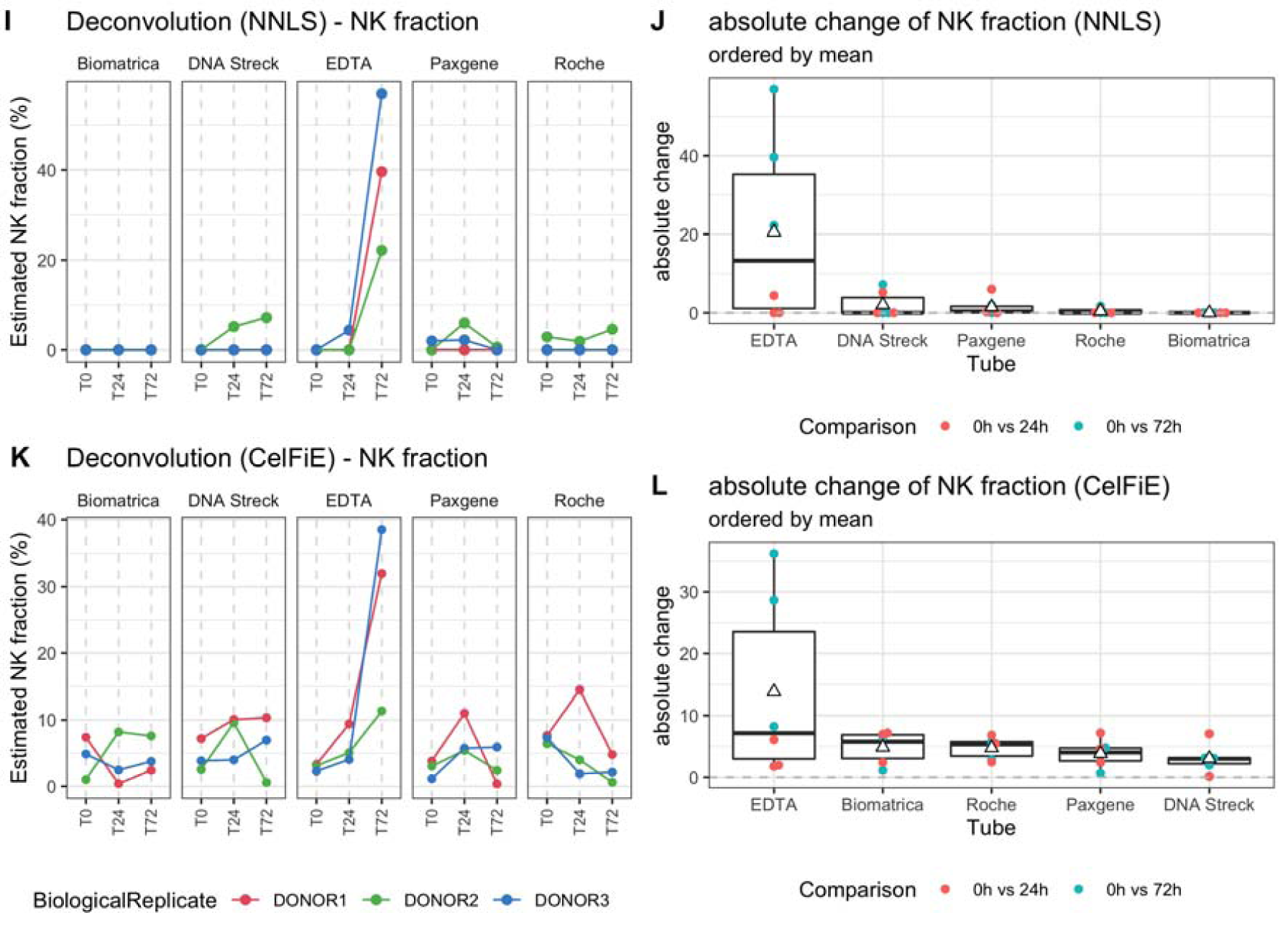
(A) Evolution of cfDNA concentration calculated based lambda DNA spike-in after DNA isolation and before library construction, (B) Boxplot of fold changes within each donor across timepoints per tube for cfDNA concentration. (C) Evolution of the genome-wide CpG methylation percentage, split per tube, (D) Boxplot of the absolute change within each donor across timepoints per tube for the genome-wide CpG methylation percentage. (E) Evolution of the absolute number of CpGs detected before setting a read count cut-off, split per tube, (F) Boxplot of the fold change within each donor across timepoints per tube absolute number of CpGs detected. (G & H) Evolution and boxplot of fold change of absolute number of CpGs detected after setting a read count cut-off of 15 reads. (I & J) Evolution of natural killer cell contribution, split per tube, calculated with NNLS using human methylation 450K and EPIC micro-array data as reference dataset or with CelFiE, using public whole genome bisulfite sequencing data from ENCODE/BLUEPRINT as reference set. (K & L): Boxplot of absolute changes in NK cell fractions within each donor across timepoints per tube. The white triangle on the boxplot corresponds to the mean of the absolute change. NK, natural killer cell. T0 = plasma prepared immediately after blood draw, T24 = plasma prepared 24 hours after blood draw, T72 = plasma prepared 72 hours after blood draw.

Overall CpG methylation (figure 3C and 3D) slightly increases in EDTA tubes across time (mean absolute change = 4.33%, median % mCpG at T0 = 43.29%, median % mCpG at T72 = 49.21%). While the absolute number of CpG covered increases in EDTA tubes across time (median number of CpGs at T0 = 3,287,720, median number of CpGs at T72 = 4,438,485), the number of CpGs with at least 15 supporting reads is relatively stable (median number of CpGs at T0 = 487,450, median number of CpGs at T72 = 517,884). In contrast to EDTA, the number of CpGs with at least 15 counts decreased in Biomatrica, DNA Streck and Roche tubes (figure 3E, 3F, 3G, 3H).

Principal component analysis reveals clustering based on the donor, indicating the blood tube, sequencing or plasma preparation protocol are not the dominant drivers in explaining the variance in the methylation pattern of the cfDNA (Figure 4). Within each donor, timepoints were evaluated for reproducibility at the CpG level. The cumulative distribution of the difference between CpG methylation fractions was visualized. On this visualization, the area left of the cumulative distribution curve (ALC) is a marker for reproducibility between these two replicates, with increasing ALC values indicating lower reproducibility (described by Mestdagh et al.^22^). The 6 ALC values per tube were similar across all tubes (supplementary figures S7, S8). Furthermore, no differentially methylated promotor regions were detected in any of the across-tube comparisons at T0 (supplementary table 2). In addition, no DMRs were observed for the Biomatrica, DNA Streck, PAXgene after comparison between T0 and T72, while the Roche tube contained 2 DMRs (MAGI1-AS1, SLX1B-SULT1A4) and the EDTA tube contained 51 DMRs (1.29%, 44 upregulated and 7 downregulated) (supplementary table 2). Among the top enriched KEGG pathways of the genes linked to the 51 DMRs, we identified upregulation of the apoptosis pathway (FDR = 0.014). FRY enrichment for GO:0006915 (apoptotic process) and GO:0030101 (natural killer cell activation) revealed significant upregulation for these pathways (FDR = 0.014 and 0.029, respectively). Further comparison between T24 and T0 in the EDTA tube revealed no DMRs.

**Figure 4:**
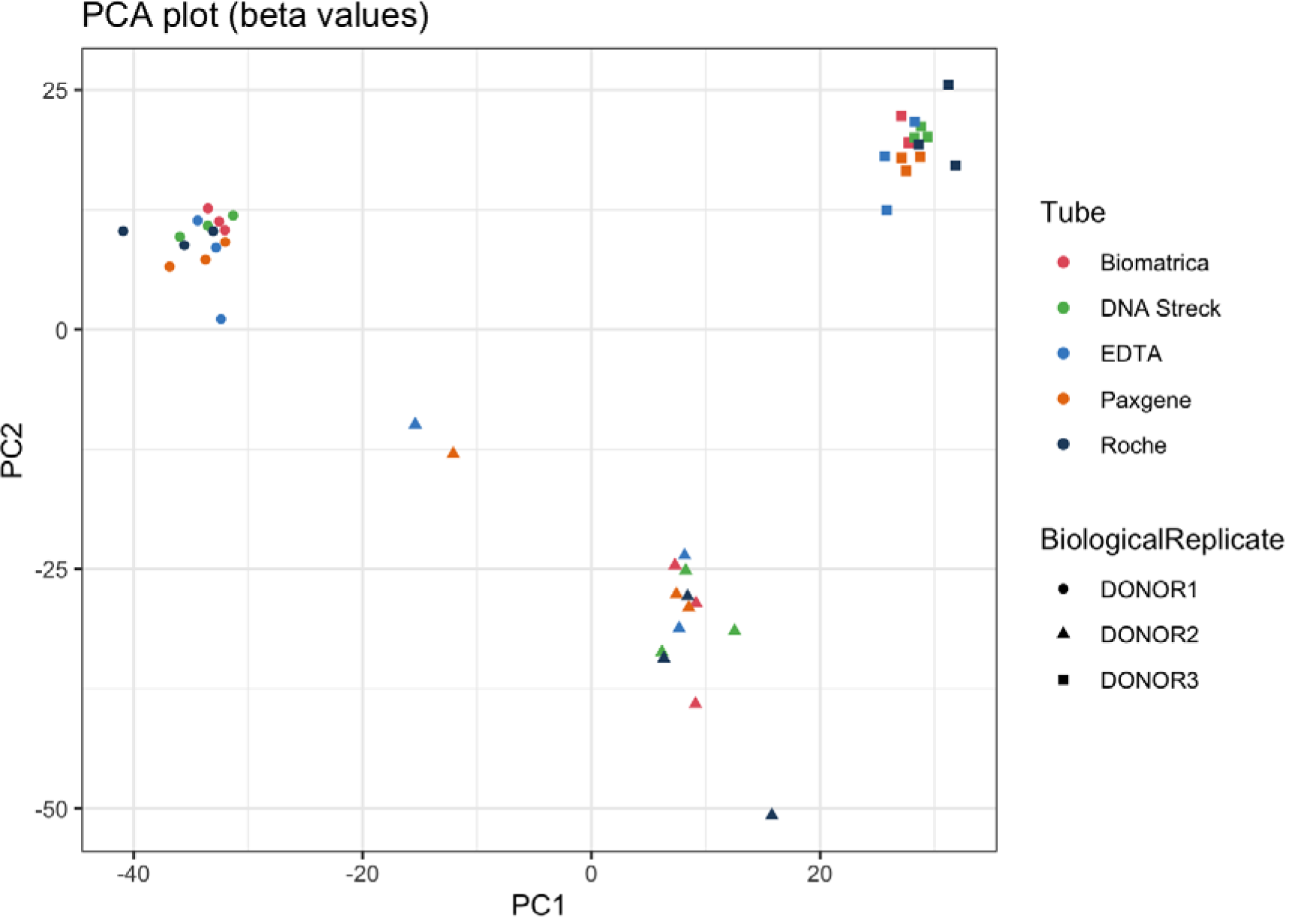
Principal component analysis performed on the beta values (fraction of methylated reads), with common cytosines in a CpG context across all samples (at least 15 reads per cytosine, n = 6706).

### Cell type contribution

In order to evaluate changes in cell contributions to cfDNA content, we performed non-negative least squares (NNLS) on the cfDNA samples in combination with a previously published cfDNA reference atlas (generated on Illumina arrays) ^3^. In order to compare cf-RRBS with Illumina HM450K or EPIC, we grouped individual CpGs into regions as described before ^2^. We observed similar proportions of erythrocyte progenitors as described by Moss et al (35.80 % [28.70-41.80]). In all EDTA tubes at T72, a sharp increase of natural killer (NK, figure 3E & 3F) cell fraction was observed (mean absolute change = 20.53, median at T0 = 0%, median at T72 = 39.6%) and a decrease of neutrophil fraction was observed (mean absolute change = 19.53, median at T0 = 40.8%, median at T72 = 7.8%, supplementary figure S9, supplementary figure S10).

In order to confirm these results, we used an independent reference dataset and deconvolution method (CelFiE) ^25^.. We observed similar changes in NK cell contribution (figure 3G & 3H), with a mean absolute change of 13.81% in EDTA tubes (median at T0 = 3.12%, median at T72 = 31.97%). The decrease of neutrophil fraction was less pronounced (supplementary figure S11, supplementary figure S12). Furthermore, CelFiE identified an overall unknown cell type percentage of 12.48% [9.36-16.59] across all samples.

## Discussion

The standardization of the pre-analytical phase is one of the major hurdles in incorporating liquid biopsy-based assays in clinical trials and routine. Several companies have developed “preservation tubes” that stabilize blood cells and prevent hemolysis and degradation of white blood cells over time. The cell-free DNA methylation pattern is currently under investigation as a marker for screening, diagnosis and therapy response monitoring^28^. We have investigated the impact of the preservation agent on the cfDNA methylation pattern in EDTA and in four preservation tubes (Roche, Streck, PAXgene and Biomatrica). Time points were selected based on a clinical situation where samples are stored at room temperature over a weekend. Although the cf-RRBS still does not investigate the full methylome, there is a strong correlation with whole genome bisulfite sequencing (r = 0.98) and SeqCap Epi CpGiant capture sequencing (Roche, r = 0.96) for the covered CpGs^17^. Thus, our results can be generalized to these bisulfite-based alternatives for cell-free DNA methylation assessment.

We were able to obtain libraries from all samples, indicating that the cf-RRBS library construction method is robust in samples with very low amounts of cfDNA (100 pg total input). Bisulfite conversion efficiency was > 99% in all tubes, indicating there is no incompatibility with the EZ DNA Lightning kit used in this study and the preservation agent (e.g. due to crosslinking of DNA). Thus, changes in genome wide CpG methylation percentage reflect changes in cfDNA composition, rather than a technical artefact.

The estimated cell-type contributions in our samples, obtained from healthy volunteers, are similar to what has previously been described by Moss^3^ and Caggiano^25^, with the majority of cfDNA is originating from leukocytes. Furthermore, a smaller but substantial fraction of the cfDNA is derived from erythrocyte progenitors in these two studies, and our results further support these findings. While earlier studies have seen that the hepatocyte-derived fraction is the second-most abundant fraction in the cfDNA (up to 10%)^6^, these more recent studies do not support this. Furthermore, our results show that the content and cfDNA yield substantially change when storing EDTA for 72 hours at room temperature. More specifically, we observed a substantial increase in NK cell contribution with both computational deconvolution and gene set enrichment analysis. Possibly, NK cells are more vulnerable to cellular stress, and degrade or lyse faster compared to other white blood cells. Previous research based on flow cytometry^29^ showed that, although NK cells are more vulnerable to storage than B or T cells, monocytes and myeloid cells are even less stable than NK cells. Our data could not confirm this. The authors recommend against performing immunophenotyping of white blood cells beyond 48 hours.

The most important limitation of this study is the small sample size. Thus, small differences should be interpreted with care. However, while only three biological replicates are available per condition (i.e. tube and time point), the quality control metrics are reproducible.

Based on the low cost of EDTA tubes, we recommend using EDTA tubes for cfDNA applications in situations where it is possible to immediately (within 24 hours) process the blood samples. If, due to logistic or practical reasons this is not feasible, a tube containing a preservation agent could be used. The type and brand of preservation tube should be selected on a per-study basis. For example, Roche tubes are not desirable in experiments that require larger amounts of DNA, as we have observed a decrease in cfDNA concentration over time. In studies where both cfRNA and cfDNA are being investigated, a preservation tube with the best overall performance should be selected^30^. For example,

DNA Streck tubes are incompatible with the analysis of extracellular RNA^31^. Other preservation tubes have been developed since the initiation of this study (e.g. Norgen cf-DNA/cf-RNA preservation tubes) and should be evaluated in a similar manner before using them for cfDNA methylation-based studies. Nikolaev and colleagues^32^ have shown that other preservation tubes mitigate the effect of other pre-analytical factors, e.g. storage temperature and long term (∼1 week) storage. While we did not investigate the effect of storage beyond 72 hours, we expect that changes in cfDNA methylation beyond 72 hours will be primarily driven by changes in cell-type contribution rather than the preservation agent used in these tubes. Still, thorough validation and quality control as emphasized by others^8,32^ remains essential. Additionally, our study adds that capillary electrophoresis with Femto PULSE is a valid approach for evaluating the presence of contaminating high molecular weight DNA, with minimal sample loss (10 pg).

In conclusion, the preservation agent used in Streck Cell-Free DNA BCT tubes, PAXgene Blood ccfDNA tubes, Roche Cell-Free DNA Collection tubes and Biomatrica LBgard blood tubes does not seem to give bias towards global or local hypo- or hypermethylation. However, EDTA tubes show a good performance in situations where they are processed within 24 hours after blood draw and thus have a higher cost/benefit ratio.

## Supplementary data

Supplementary table 1: sample characteristics.

Supplementary table 2: differentially methylated regions

Supplementary figures: additional quality control figures, full deconvolution results and additional Femto PULSE figures.

## Data availability

Scripts, supporting files and full data analysis is available at https://github.com/rmvpaeme/cfRRBS_tube_study.

Raw data is available at EGA identifier EGAD00001006007. Processed data can be found at ArrayExpress identifier E-MTAB-8858.

The cf-RRBS protocol is available at https://www.protocols.io/private/4098389D37C151C92AA19EC574AD9201.

## Acknowledgements

We thank all healthy volunteers for participating in this study. A.D.K. and R.V.P. were funded by a predoctoral fellowship from the Research Foundation Flanders (FWO). B.D.W is a senior clinical investigator for the FWO. This project was partially funded by Kom op tegen Kanker, a CRIG (Cancer Research Institute Ghent) Young Investigator Proof-of-Concept grant (to B.D.W.), and by the VIB Grand Challenges Program (Nico Callewaert lab). We thank Peter Stockwell for assistance with mkrrgenome. We thank Charlotte Vandeputte, Eveline Debals, Malaϊka Van Der Linden, Thalia Van Laethem, Eveline Vanden Eynde, Kimberly Verniers, Peter Degrave, Dimitri Broucke, Machteld Baetens, Aline Eggermont and Els De Smet for assistance with plasma preparation, cfDNA extraction, library preparation and sequencing.

We thank the members of the extracellular RNA quality control consortium (exRNAQC) at Ghent University (Eva Hulstaert, Hetty Helsmoortel, Kimberly Verniers, Francisco Avila Cobos, Anneleen Decock, Eveline Vanden Eynde, Céleste Van der Schueren, Jasper Anckaert, Justine Nuytens, Jill Deleu, Nurten Yigit, Celine Everaert, Jilke De Wilde, Kathleen Schoofs, Gary Schroth, Scott Kuersten, Nele Nijs, Carolina Fierro, Olivier De Wever, An Hendrix, Bert Dhondt, Annouck Philippron) for assistance with experimental setup, data-analysis and interpretation. The computational resources (Stevin Supercomputer Infrastructure) and services used in this work were provided by the VSC (Flemish Supercomputer Center), funded by Ghent University, FWO and the Flemish Government – department EWI.

## Disclosures

A.D.K is listed as co-inventor in patent application PCT/EP2017/056850 related to the cf-RRBS method. Roche provided the cell-free DNA collection tubes free of charge. Roche was not involved in study design or writing of this manuscript.

## Notes

https://github.com/rmvpaeme/cfRRBS_tube_study

https://github.com/rmvpaeme/cfRRBS_manuscript

https://www.ebi.ac.uk/arrayexpress/experiments/E-MTAB-8858/

### Summary of Updates

a reader notified us that "One study suggests that PAXgene Blood ccfDNA tubes and Streck Cell-Free DNA BCT tubes are unsuitable for the analysis of methylated cfDNA, as assessed with methylation-specific PCR (MSP) for SEPT9" should read "One study suggests that PAXgene Blood DNA tubes and Streck Cell-Free DNA BCT tubes are unsuitable for the analysis of methylated cfDNA, as assessed with methylation-specific PCR (MSP) for SEPT9"

