## supplemental figures for "Genome-wide study of the effect of blood collection tubes on the cell-free DNA methylome"

### Blood collection preservation tubes have limited effect on methylation status of cell-free DNA: supplementary figures

#### Code and data availability

The full data-analysis, including markdown file with these (updated) figures and raw data values are available at <https://github.com/rmvpaeme/cfRRBS_tube_study>.

#### Sequencing and preprocessing QC

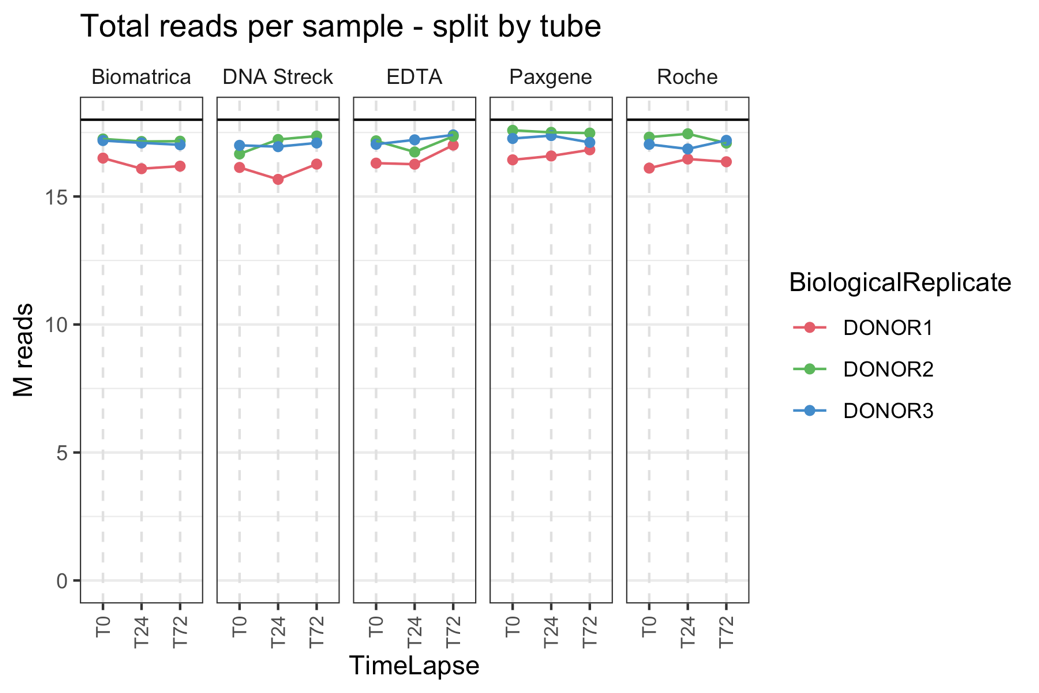

Figure S1: total reads remaining after subsampling and adaptor trimming with trim galore. Solid black line indicates 18M reads (= level to which all libraries were subsampled).

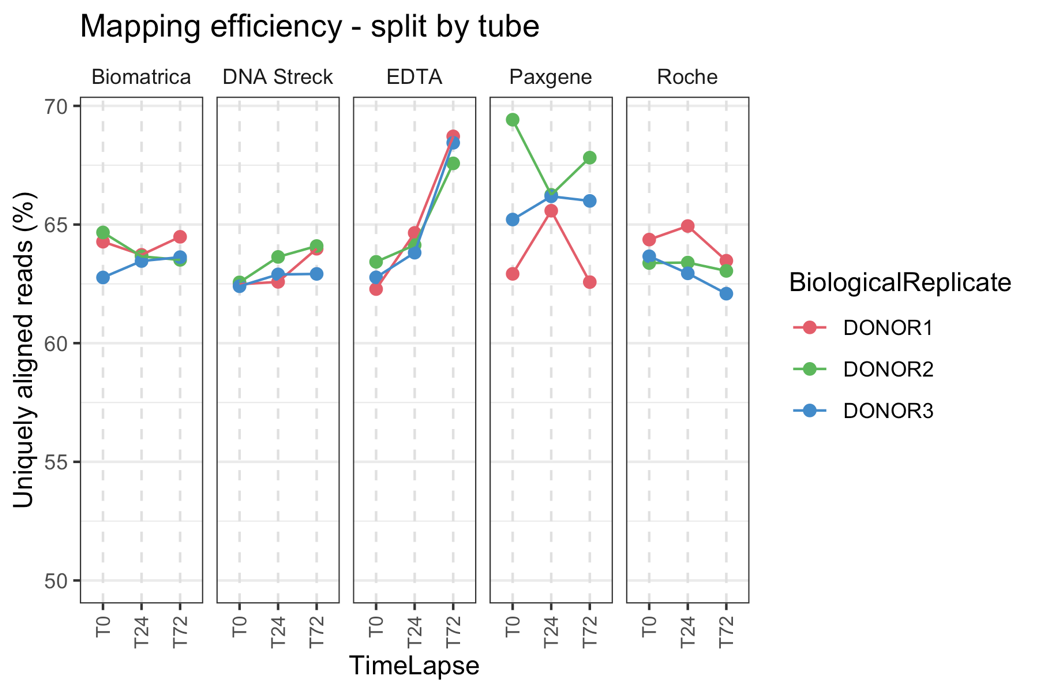

Figure S2: percentage of reads mapping uniquely mapping to GRCh37, as evaluated with Bismark.

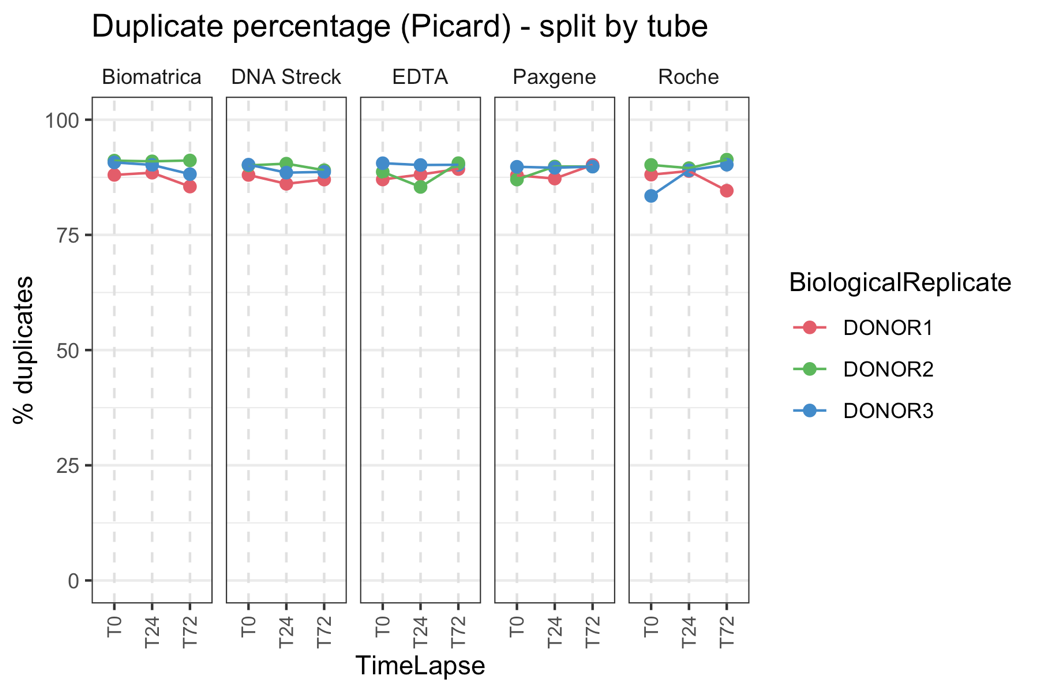

Figure S3: duplicate percentage after subsampling and removal of optical duplicates, calculated by Picard MarkDuplicates.

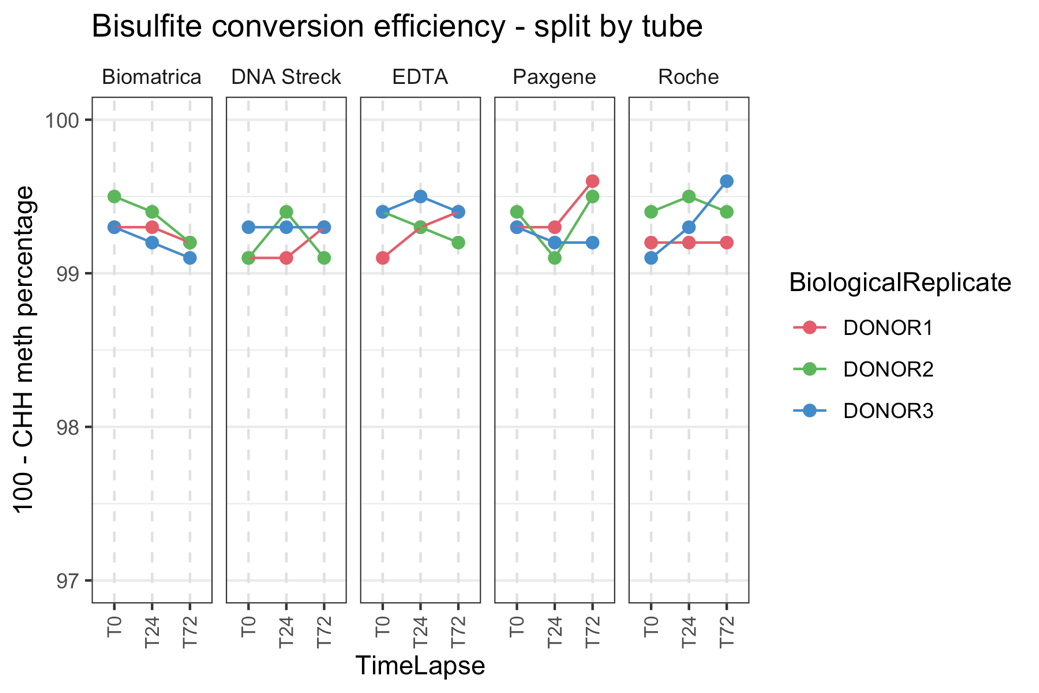

Figure S4: bisulfite conversion efficiency evaluated by CHH methylation status. Theoretically, (100 - CHH methylation %) should be 100, as this equals an 100% conversion efficiency.

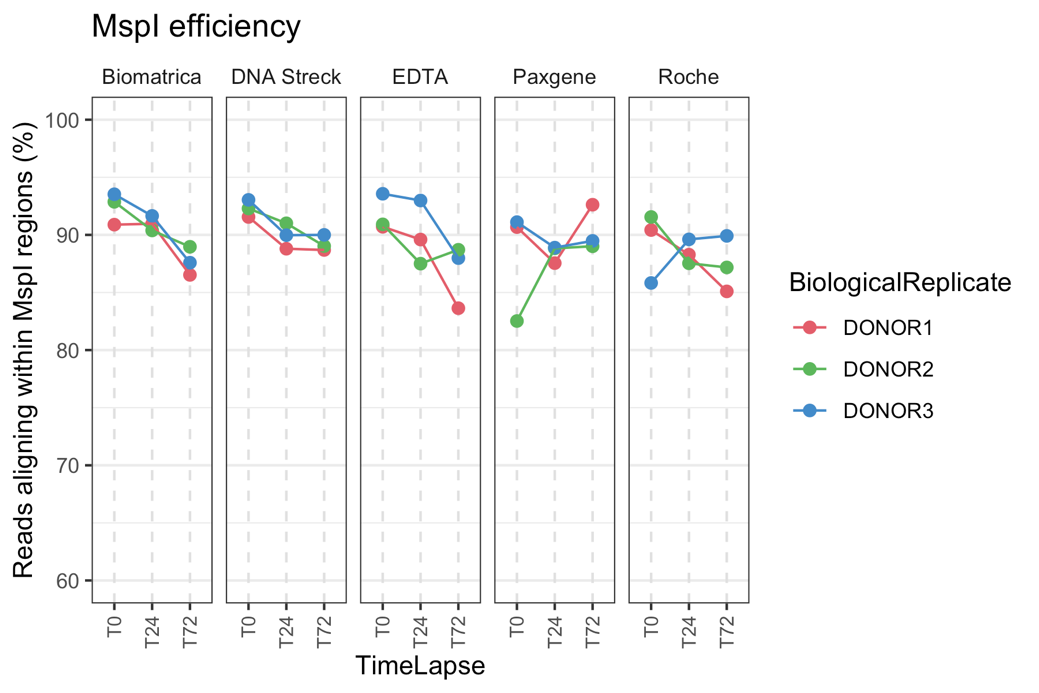

Figure S5: MspI efficiency, evaluated by calculating the percentage of reads mapping within MspI regions between 20-200 bp (see methods of main manuscript).

#### Tube content

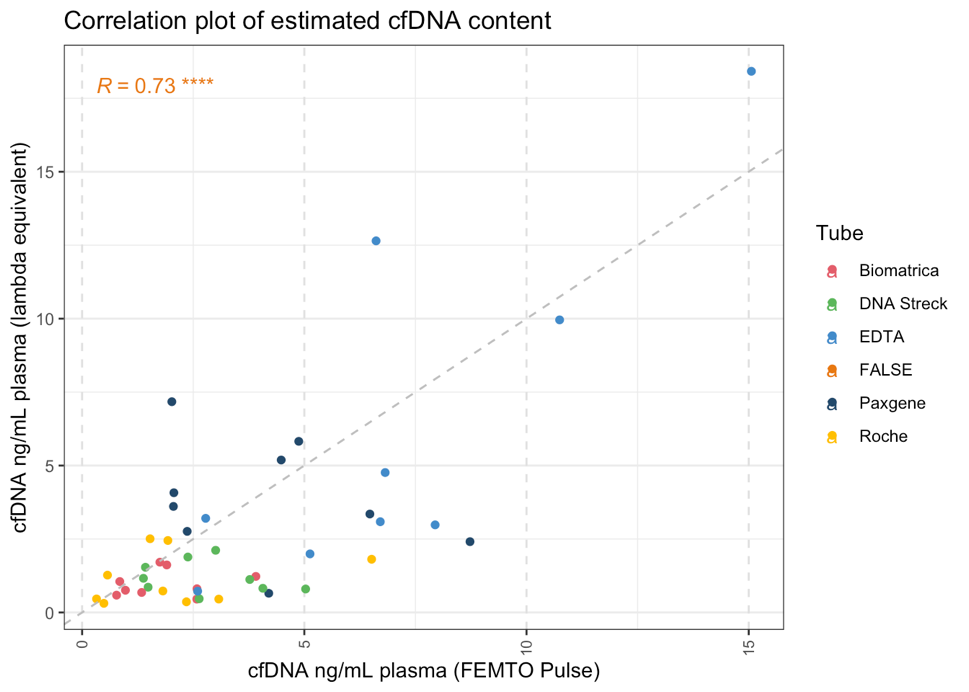

Figure S6: Scatter plot and Pearson correlation of cfDNA content estimated with capillary electrophoresis and lambda DNA spike-in, in ng/mL plasma.

#### Reproducibility

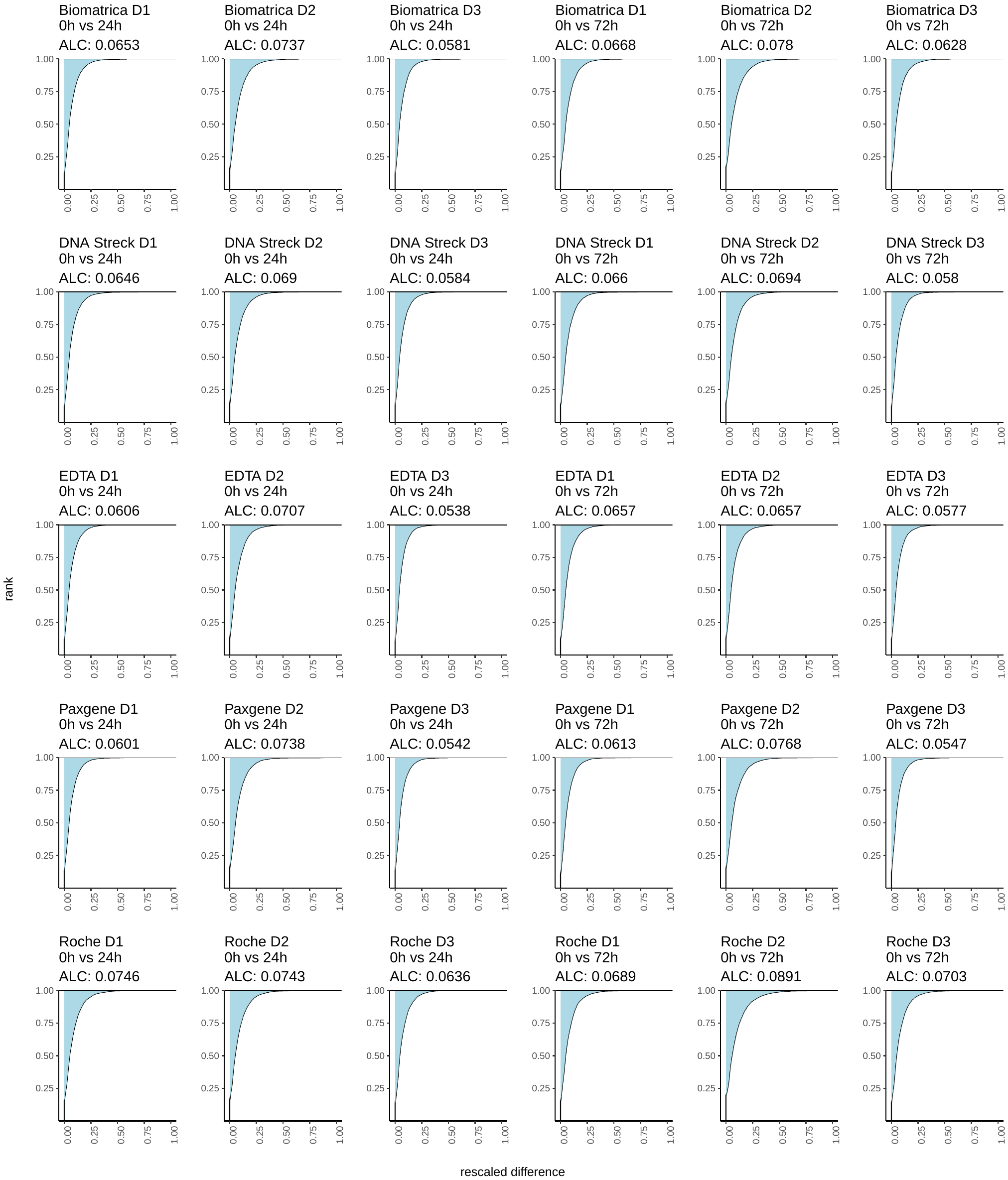

Figure S7: individual cumulative distribution plots, illustrating the area-left-of-the-curve (ALC, blue shaded area, see Mestdagh et al.^1^), a marker for reproducibility between two replicates. Increasing values indicate lower reproducibility between two conditions.

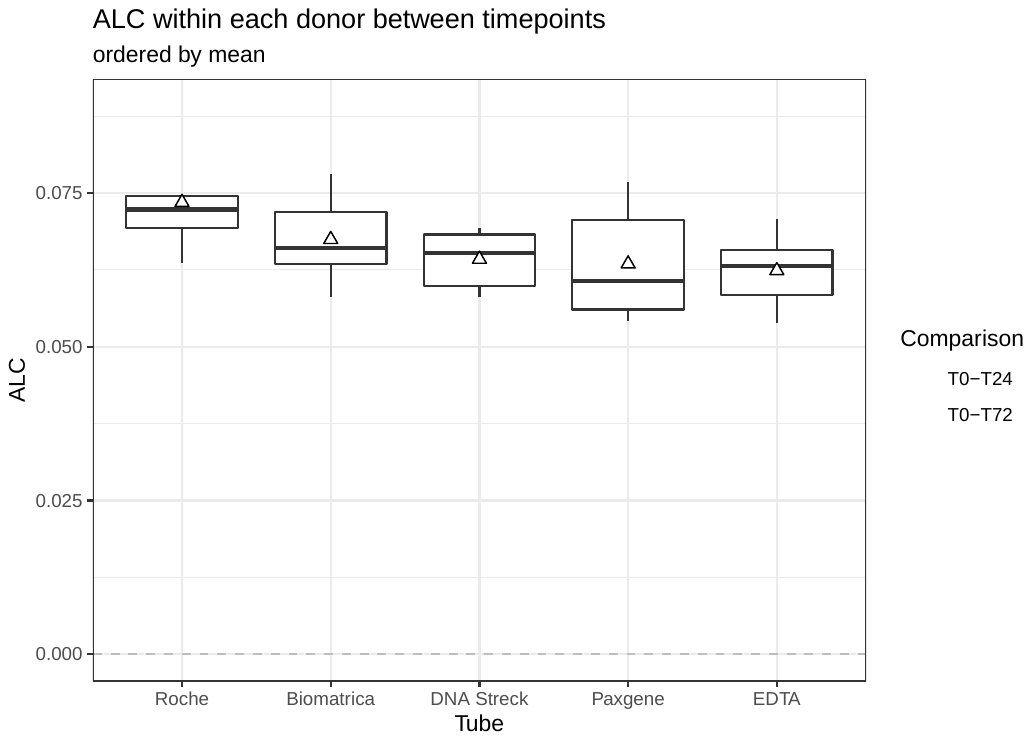

Figure S8: reproducibility assessed by area left of the curve. In order to assess reproducibility per individual CpG methylation status between timepoints within one donor, we calculated the area between the theoretical cumulative distribution representing perfect reproducibility (optimal curve) and the cumulative distribution of the timepoint-comparison. This area left of the cumulative distribution curve (ALC) decreases with increasing reproducibility and is approximated by the mean absolute replicate expression difference (described by Mestdagh et al.^1^).

#### NNLS deconvolution

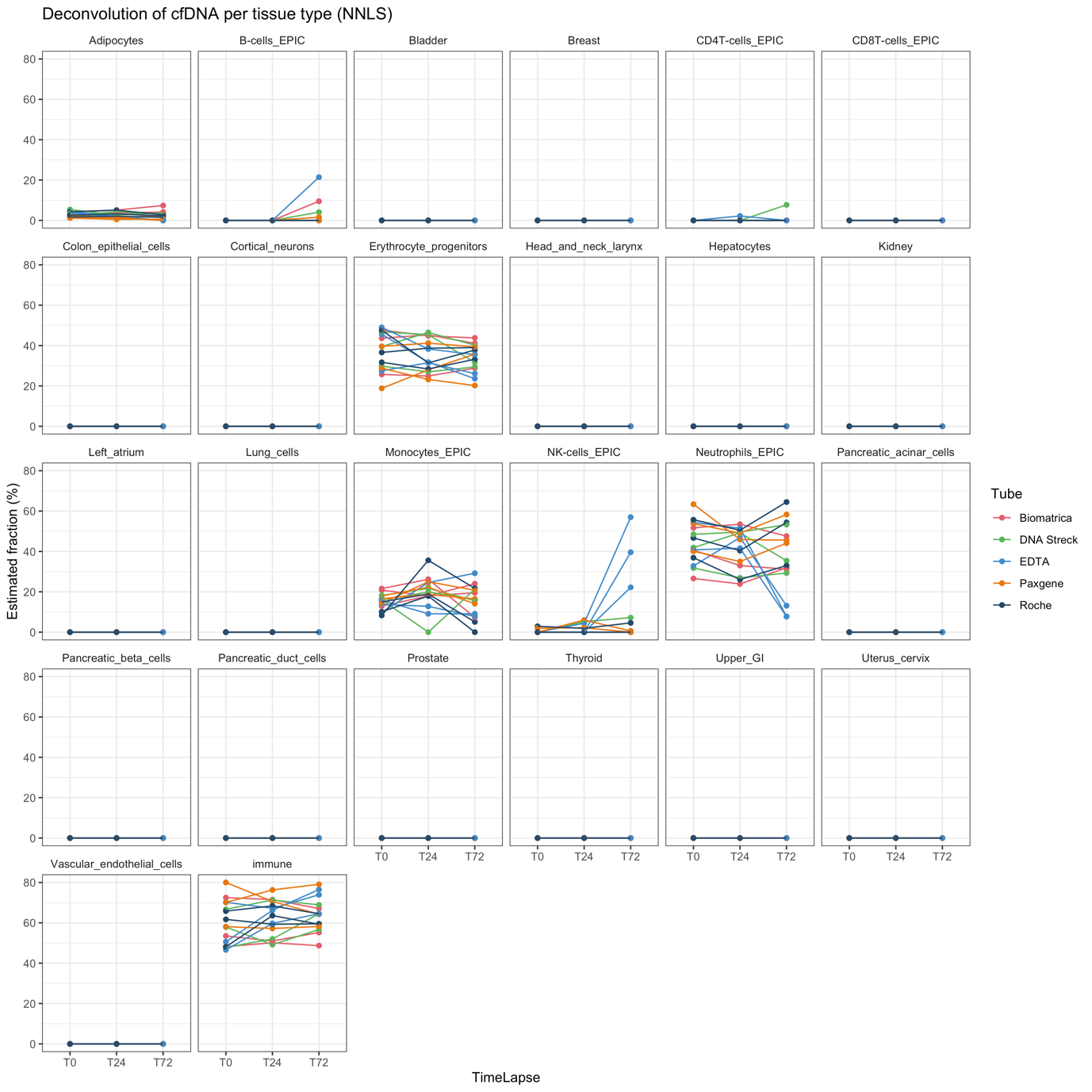

Figure S9: proportions split by tissue type with NNLS, using the data from Moss et al.^2^ as reference set. “Immune” refers to the sum of B-cells, T-cells, monocytes, NK-cells and neutrophils.

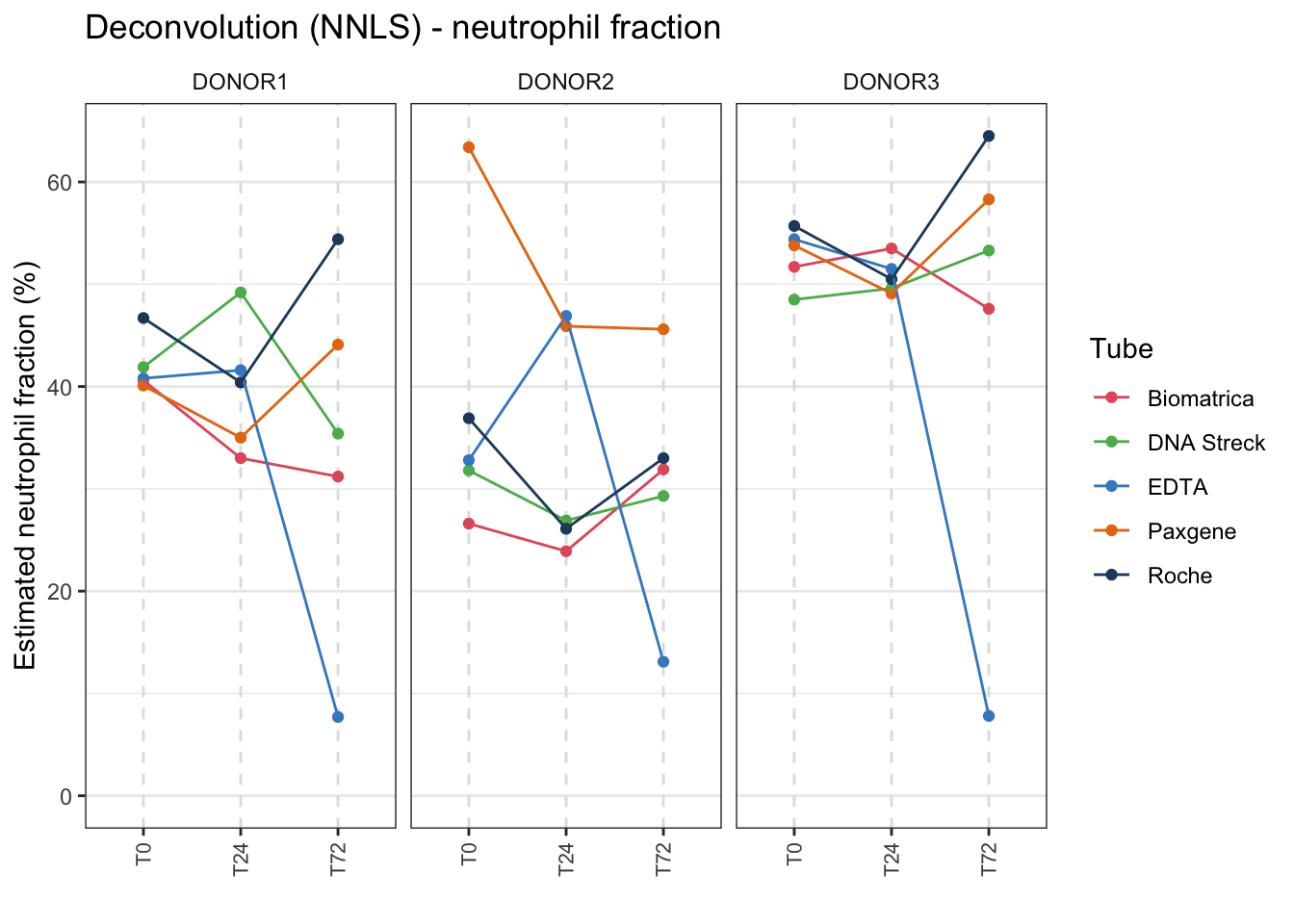

Figure S10: estimated neutrophil fraction with NNLS, using the data from Moss et al.^2^ as reference set.

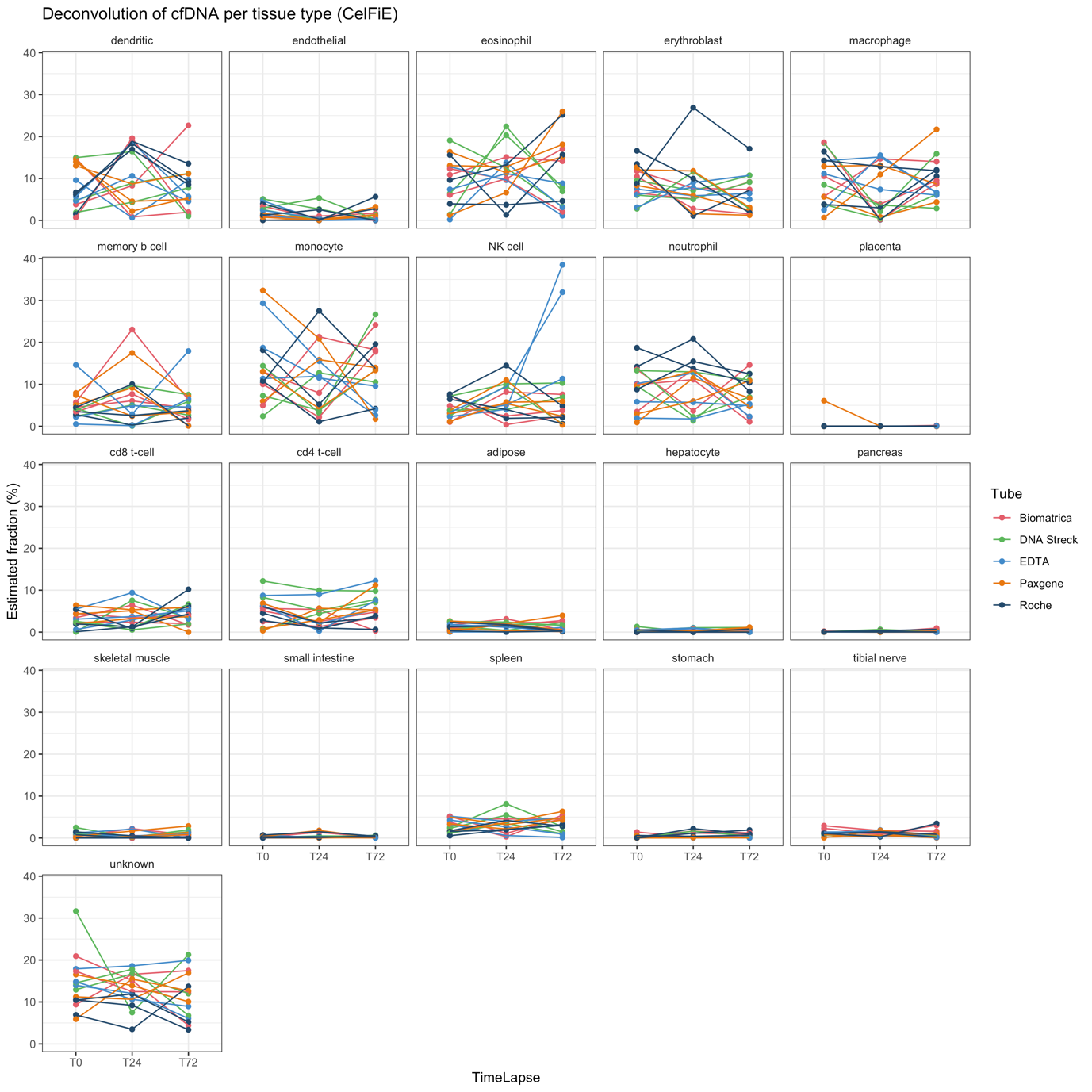

Figure S11: proportions split by tissue type with CelFiE, using the data from Caggiano/ENCODE/BLUEPRINT^3^ as reference dataset.

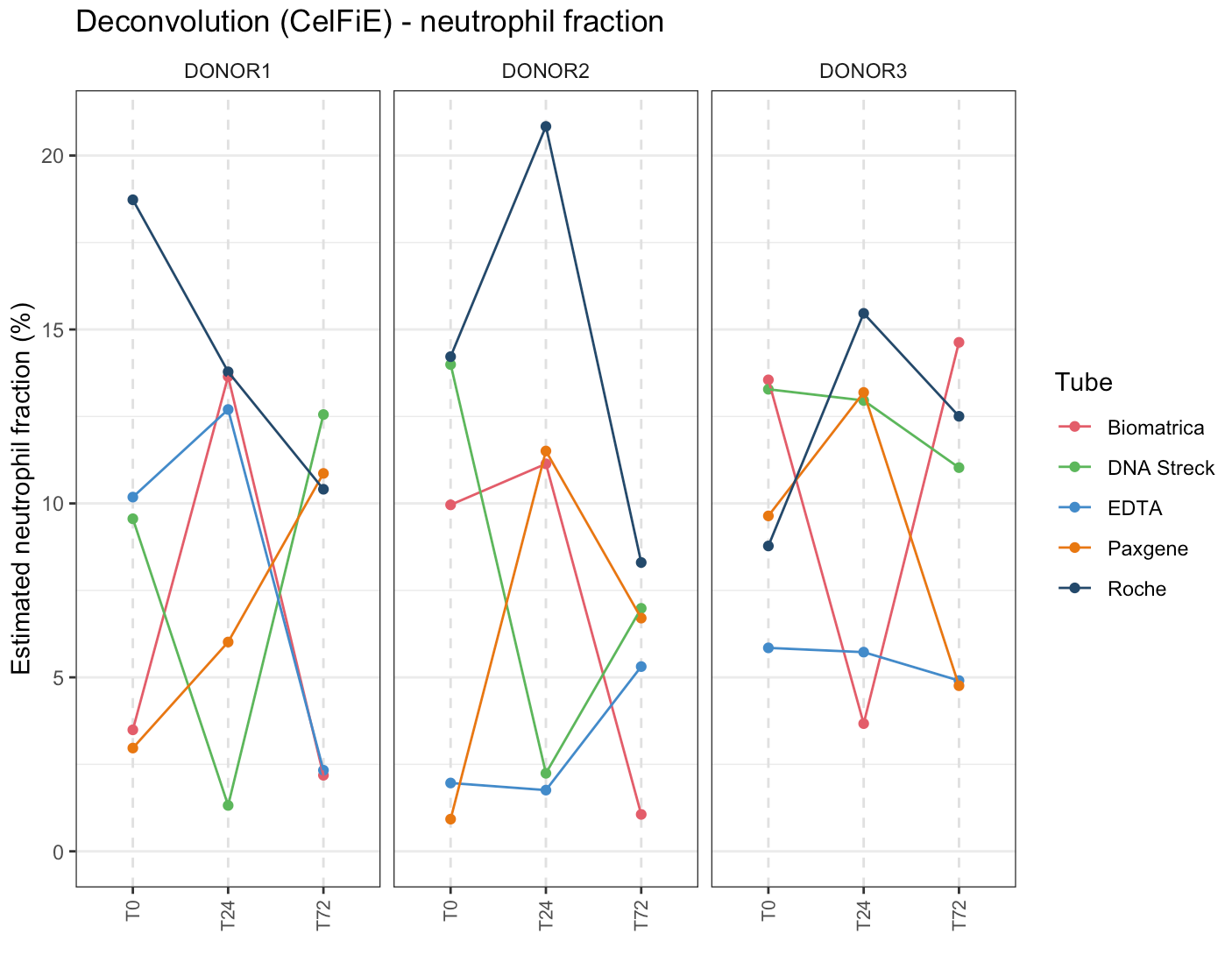

Figure S12: estimated neutrophil fraction with CelFiE, using the data from Caggiano/ENCODE/BLUEPRINT^3^ as reference datatset.

#### Femto PULSE

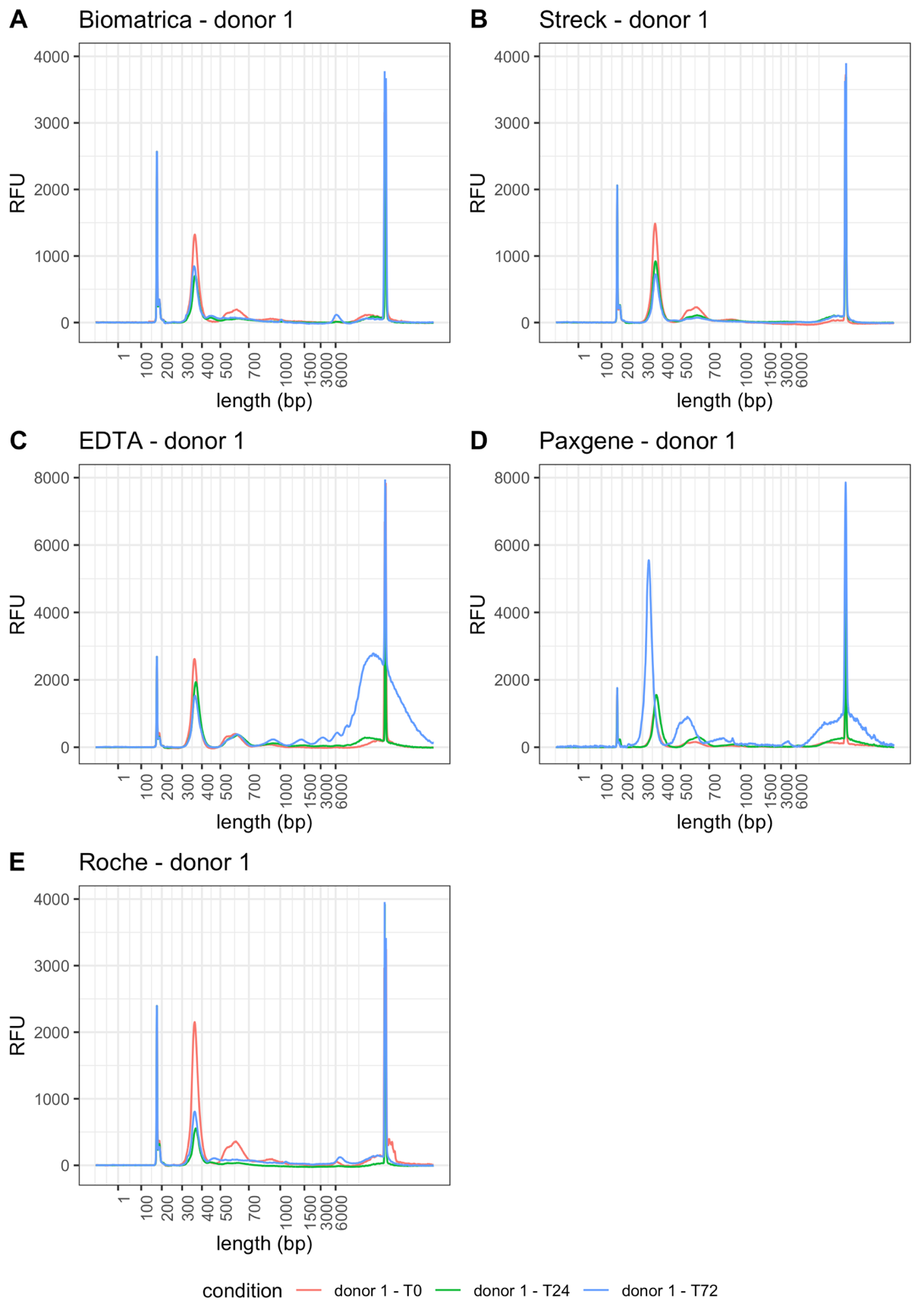

Figure S13: Capillary electropherograms (Femto PULSE) of all 15 cfDNA samples obtained from donor 1. RFU = relative fluorescence units. T0 = plasma prepared immediately after blood draw, T24 = plasma prepared 24 hours after blood draw, T72 = plasma prepared 72 hours after blood draw. Peaks at 1 and 6000 bp correspond to the lower marker (1 bp) and upper marker (6000 bp).

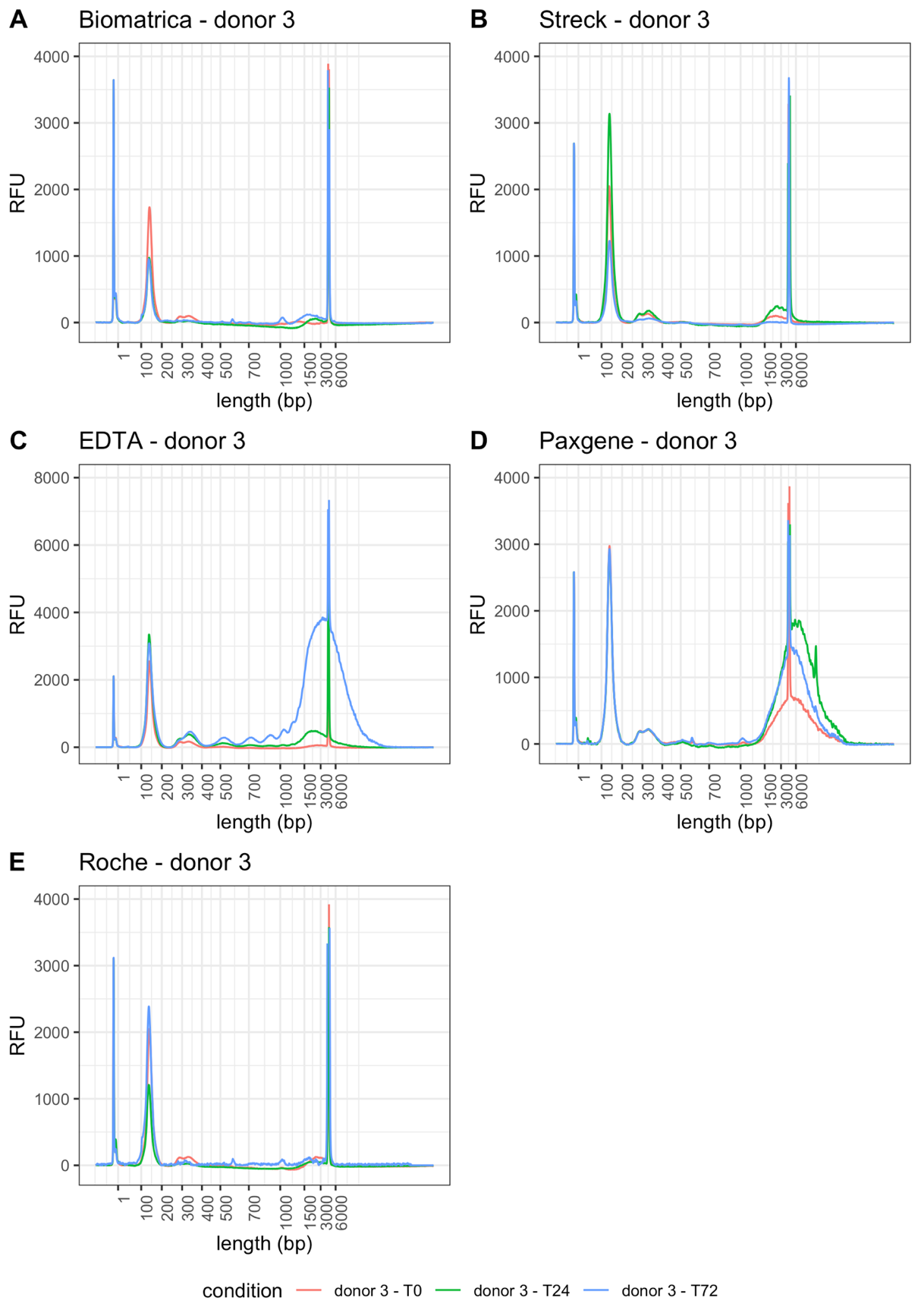

Figure S14: Capillary electropherograms (Femto PULSE) of all 15 cfDNA samples obtained from donor 3. RFU = relative fluorescence units. T0 = plasma prepared immediately after blood draw, T24 = plasma prepared 24 hours after blood draw, T72 = plasma prepared 72 hours after blood draw. Peaks at 1 and 6000 bp correspond to the lower marker (1 bp) and upper marker (6000 bp).
